# AlliGator: Open Source Fluorescence Lifetime Imaging Analysis in G

**DOI:** 10.1101/2025.05.22.655640

**Authors:** X. Michalet

## Abstract

Fluorescence Lifetime Imaging (FLI) is a technique recording the temporal decay of fluorescence emission at every pixel of an image. Analyzing the information embedded in FLI dataset requires either fitting the decay to a predefined model using nonlinear least-square fit or maximum likelihood estimation, or projecting the decay on an orthogonal basis of periodic functions as in phasor analysis. AlliGator is a Windows open source software (BSD license) implementing these approaches in a user-friendly graphical user interface (GUI) and offering numerous unique features such as Förster Resonant Energy Transfer (FRET) stoichiometry analysis, or the creation of maps of lifetime-derived quantities such as membrane potential. It leverages the unique ability of the LabVIEW graphical programming language (G) to design feature-rich GUI, and supports user-developed plugins written in python to extend its native capabilities.

## Metadata

### 1. Motivation and significance

Fluorescence lifetime imaging (FLI) techniques have gained in popularity in the biomedical field thanks to technological advances in laser sources and detectors, both in its microscopic version (FLI microscopy or FLIM) [1] and its mesoscopic or macroscopic version (mFLI or MFLI) [2, 3, 4]. However, FLI data analysis remains challenging as different analytical approaches are available, such as nonlinear least-square fit (NLSF), maximum likelihood estimation (MLE) [5], phasor analysis [6] or deep learning [7], among others, each requiring expertise to implement, and once data is processed, to interpret.

**Table 1:** Code metadata (mandatory)

| **Nr.** | **Code metadata description** | **Metadata** |
| --- | --- | --- |
| C1 | Current code version | 1.01 |
| C2 | Permanent link to code/repository used for this code version | https://github.com/smxplorer/alligator |
| C3 | Permanent link to Reproducible Capsule | https://doi.org/10.5281/zenodo.15686834 |
| C4 | Legal Code License | BSD |
| C5 | Code versioning system used | git |
| C6 | Software code languages, tools, and services used | LabVIEW 2021 & Python 3.9-3.10 |
| C7 | Compilation requirements, operating environments & dependencies | LabVIEW 2021 SP1 64-bit, Vision Development Module 2021, OpenG Toolkit, etc. |
| C8 | If available Link to developer documentation/manual | https://github.com/smXplorer/AlliGator/tree/main/src-docs |
| C9 | support email |  |

**Table 2:** Software metadata (optional)

| **Nr.** | **(Executable) software meta-data description** | **Please fill in this column** |
| --- | --- | --- |
| S1 | Current software version | 1.01 |
| S2 | Permanent link to executables of this version | https://github.com/smXplorer/AlliGator/tree/main/installer |
| S3 | Permanent link to Reproducible Capsule | https://doi.org/10.5281/zenodo.15686834 |
| S4 | Legal Software License | BSD |
| S5 | Computing platforms/Operating Systems | Microsoft Windows |
| S6 | Installation requirements & dependencies | LabVIEW 2021 SP1 64-bit Runtime Executable, Vision Development Module 2021 Runtime Executable |
| S7 | link to user manual | https://alligator-distribution.readthedocs.io |
| S8 | Support email for questions |  |

AlliGator was developed with the aim of providing maximum flexibility to explore the effect of changing parameters on the results of different analysis approaches, while keeping the interface purely graphical and interactive. It has been used in scientific publications covering techniques as diverse as wide-field m/MFLI using an intensified CCD camera [8, 9], a linear SPAD array [10] and a SPAD camera [11, 12], or FLIM using time-resolved confocal microscopy [13] or a SPAD camera [14], and is continuously being improved with additional features.

The software takes advantage of the ease of developing a feature-rich graphical user interface (GUI) using the LabVIEW graphical programming environment (LabVIEW is a trademark of National Instruments) and its image handling library (Vision Development Module or VDM). AlliGator distinguishes itself from a number of similar open source projects (a long and fairly comprehensive list of which can be found on the documentation page of the PhasorPy project [15] reproduced in part in Appendix A) by supporting datasets requiring per-pixel instrument response function or phasor calibration (as is required with SPAD cameras), as well as by numerous ways of analyzing data from single or multiple ROIs, both in terms of fitting methods and the phasor approach. While its core is developed using LabVIEW, a simple API to support Python plugins is included, which does not require any access to the LabVIEW source code or compiler. A Windows installer for a standalone version of the software is released with each new version, that only requires a license for the VDM runtime.

### 2. Software description

The software comprises a main window, which provides different visualizations of the dataset under analysis (left side of Fig. 1) as well as as different analysis outputs in linked panels (right side of Fig. 1). Accessory windows allow the user to set analysis parameters, explore the dataset at the single-pixel or single-region of interest (ROI) level, perform line profile analysis or manipulate and fit histograms of multiple analysis outputs. The user interacts with the GUI using buttons, window menus and graphical objects’ contextual menus, as well as via mouse-based actions on images, graphs, slides and most other GUI objects.

**Figure 1.**
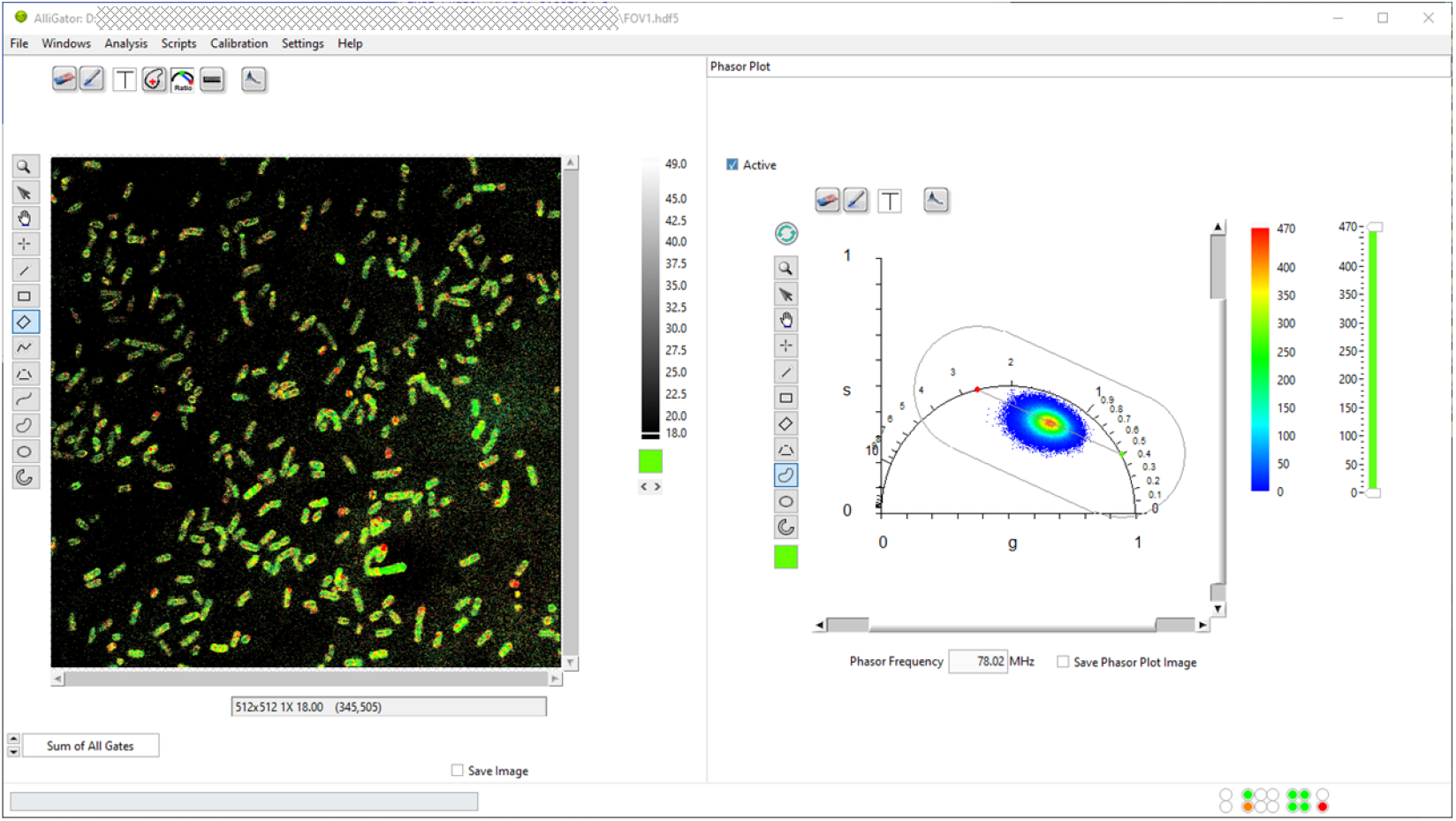
AlliGator’s main window during the final stage of an example of FLI analysis described in section 3. The left side of the window shows a dataset after overlay of the color-coded amplitude-averaged lifetime. The right side of the window show the `Phasor Plot` panel in which the pixel-wise phasor plot is represented. Two references (green and red dots) used for linear decomposition of each phasor are also represented, together with the segment connecting them and the region within which linear decomposition analysis is confined.

#### 2.1. Software architecture

AlliGator is built around a simple producer-consumer architecture. The main producer is the user-event loop hosted by the main window (see Fig. 1 for the UI and Fig. 2 for an overview of the software architecture), which registers all user interface interactions and translates them into action requests added to the main AlliGator action queue (green Q icon in Fig.2).

**Figure 2.**
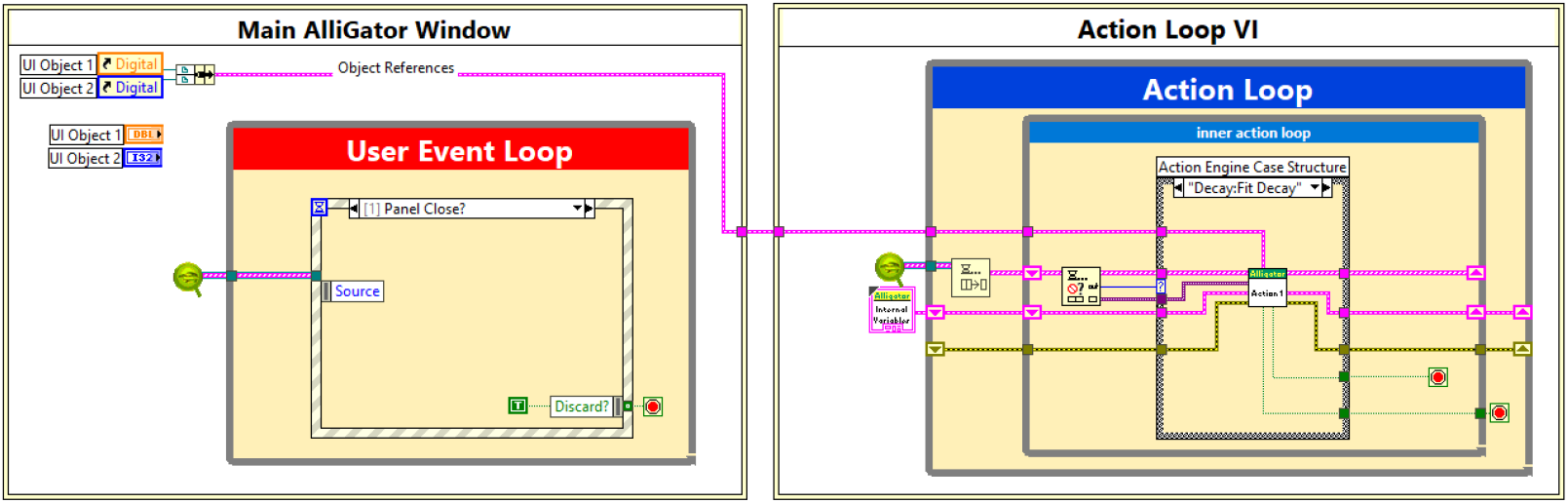
Simplified architecture of the AlliGator software. The `Main AlliGator Window` virtual instrument (VI), whose simplified code is shown to the left, contains a user-event loop which registers user interactions with various UI objects and passes corresponding action requests via a queue (green Q icon) to the `Action Loop VI` (shown on the right) running in parallel. Ancillary windows opened by the `Main AlliGator Window` (not shown) also send action requests to the Action Loop VI. The action loop handles *scripts* (arrays of action requests) in a first-in first-out order, and can insert new action requests within a script. Actions are processed by a hierarchy of sub-VIs organized into specialized libraries (e.g. file loading, phasor analysis, NLSF, etc.). Update requests to ancillary windows are handled via custom user-events. Note that in the actual code, the two pieces of code shown in the two boxes are disconnected (run in parallel), the references to the main window’s objects being passed dynamically at launch time to the `Action Loop VI`. A hierarchical diagram of all VI libraries comprising the software is shown in Fig. B.5.

The consumer is a finite state machine loop (shown on the right of Fig. 2) running in a separate routine (or *Virtual Instrument*, as an individual piece of LabVIEW graphical code is called, or *VI* for short), which dequeues action requests and processes them in a first-in first-out order.

Ancillary windows (`Histogram, Settings, Image Profile, Local Decay Graph`, etc.) launched from the main AlliGator window by user actions such as selection of window menu items or right-click on contextual menu items, behave in the same manner, sending action requests via the main queue, and receiving UI update data via dynamic events.

Internal data, including the current dataset and results of different analysis steps, are stored in a single structure passed along as a reference to sub-VIs involved in data processing, avoiding unnecessary data copies. User-defined settings can be interactively changed via a dedicated multi-panel `Settings` window, these settings being accessible (and modifiable) anywhere in the code.

Thanks to the built-in parallelism of LabVIEW code, a number of computational tasks such as NLSF calculations can take advantage of the computer’s multicore architecture and multithreading capabilities, speeding up analysis of large datasets.

#### 2.2. Software functionalities

AlliGator supports datasets files from major FLI data acquisition hardware, including confocal microscopes or intensified CCD (ICCD) and wide-field SPAD cameras. These datasets can consist of either streams of photons with position, macrotime and microtime information for each photon, or a series of images, each image corresponding to a finite temporal interval within the laser period, from which the fluorescence decay at each pixel can be reconstructed. It also allows saving datasets into a highly efficient and open source file format based on the Hierarchical Data Format (HDF5) (see the *AlliGator HDF5 File Format* online manual page for details [16]). Analysis results can be saved as text files (e.g. comma-separated-values or CSV) for further processing by third party software, or reloaded later for processing with AlliGa-tor. Information pertaining to ROIs, phasor calibration, phasor references, or detailed phasor or NSLF analysis outputs, are saved in easy to read eXtended Markup Language (XML), Javascript Object Notation (JSON) or HDF5 file formats, which can be reloaded for reproducibility purposes. Datasets can be averaged, binned, background- or pile-up-corrected when applicable, and single-pixel or region-of-interest (ROI) decays can be processed in a variety of optional ways easily configurable via the `Settings` window.

AlliGator supports multiple user-defined ROI shapes as well as algorithmically defined ones (grids, ROIs with pixel intensities over a threshold, image mask, etc.), which can be complemented by user-defined Python plugins if needed. ROIs can be merged or on the contrary, decomposed into individual pixel ROIs. Single or multiple ROIs, as well as all pixels of an image, can be analyzed using single- or double-exponential fits using weighted or unweighted NLSF or MLE, as well as by phasor analysis. Maps of fit parameters (lifetimes, amplitudes, etc.) or phasor-derived quantities (fraction of a 2-component mixture, average lifetime, etc.) can be overlaid as color-coded maps on the source image, exported as images or CSV matrix files, or further processed to extract statistical histograms or scatter-plots of one parameter versus another, among other analysis workflows described in the online manual [16]. These operations are accessible via right-click menus in the respective graphs objects representing the results of different analyses. Histograms of parameters can be fitted with (multi)peak models or exported as CSV files for external processing.

Similarly, phasor analysis can be performed at the single- or multiple-ROI, or whole-image level. ROIs can be defined in the phasor plot in order to highlight the corresponding regions of the source image, or to limit data analysis to a subset of the computed phasors. AlliGator offers different ways to define the so-called phasor calibration at the core of any phasor analysis, and allows saving calibration to file for further reuse. It also provides a collection of phasor processing functions.

Most importantly, every single action and analysis output is automatically exported into a Rich Text Format (RTF) notebook, in which the user can also copy and paste any image or graph from AlliGator’s many windows, or from external sources, as well as type any text needed to further document an analysis workflow.

Finally, user plugins can be developed in Python to add new analysis functions by taking advantage of the simple application programming interface (API) described in the corresponding section of the manual [16]. A Python plugins folder is used to store Python scripts containing normal Python functions with a few additional commented sections instructing AlliGator on (i) where to insert the function as a menu item, (ii) what data to pass to the function, (iii) what data output to expect from the function and (iv) where to display that data. Several examples are provided with each release.

#### 2.3. Sample code snippet analysis

Fig. 3 shows a graphical code snippet (or diagram) corresponding to one example of data analysis in AlliGator. This particular analysis determines the intersection of the principal axis of inertia of the phasors with the Universal Circle (UC), that is, the locus of single-exponential decays in the phasor plot [6]. The corresponding two phasors (corresponding to two single-exponential decays and referred to as *phasor references*) can then be used for linear decomposition of any phasor into a weighted sum of these references.

**Figure 3.**
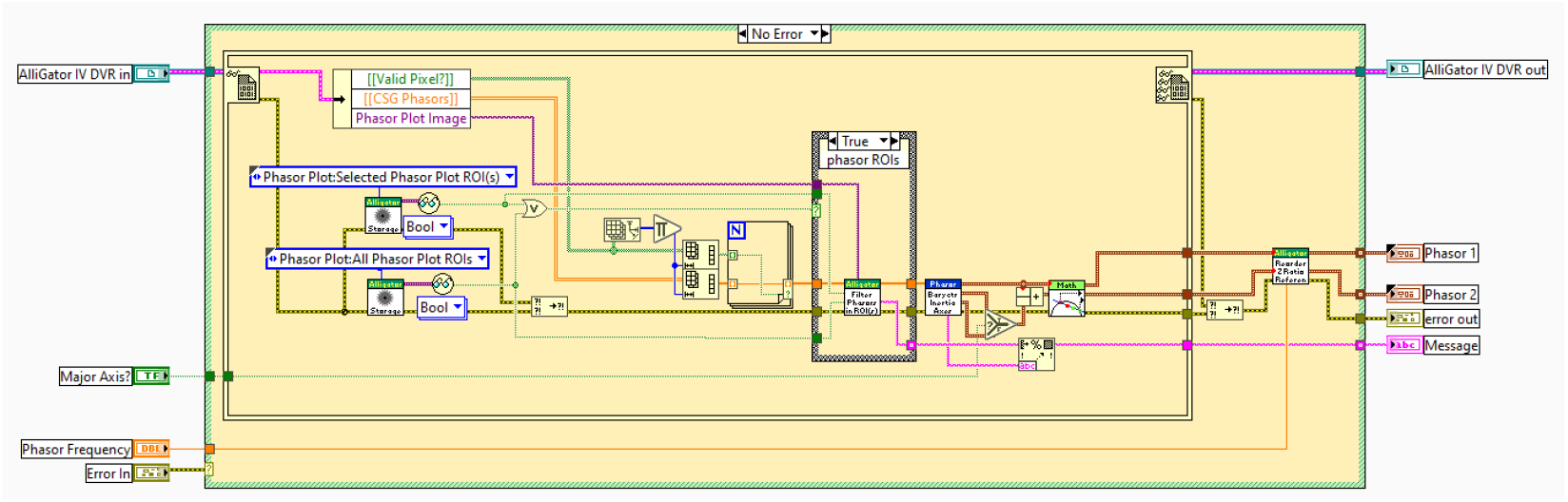
Example of AlliGator code: `AlliGator Use Phasor Plot UC-Principal Axis Intersections as References .vi.` This snippet computes the minor or major axis of the phasor plot and determines its intersection with the universal circle (UC), the locus of single-exponential decays in the phasor plot, and uses the two corresponding phasors as references for subsequent mixture analyses.

This VI takes as inputs the internal variables’ data value reference (`AlliGator IV DVR`), the `phasor frequency` as well as a boolean flag (`Major Axis?`) specifying which principal axis to use for the analysis. It then retrieves the necessary internal data (2D array of `Valid Pixels?` flags, 2D array `CSG` Phasors of single-precision complex phasor values), and user-defined settings defining whether to limit the analysis to one or more of the ROIs defined in the phasor plot image, and computes the major and minor axis before determining the intersection of the selected axis with the UC. The resulting intersections (`Phasor 1 and Phasor 2`) are then output together with an optional string (`Message`) describing the phasor ROIs used in the analysis. Subsequent VIs (not shown) called after this code snippet display these phasor references in the phasor plot image and graph (e.g. Figs. 1 & 4c), and save them internally. This code snippet illustrates the visual simplicity of properly laid out graphical (G) code, each square icon with a banner in the diagram representing a sub-VI (sub-routine), whose graphical code can be accessed by double-clicking on it in the LabVIEW development environment. This process can be repeated until only primitive functions are involved in the sub-VI’s code.

The full list of VIs comprising AlliGator and a brief description of their function can be found in the src-docs folder of the source code repository [17].

### 3. Illustrative example: Bacterial membrane potential analysis using voltage sensitive dyes

Here, we briefly provide an overview of the type of analysis used in a recent article studying the membrane potential of bacteria using voltage sensitive fluorophores whose lifetime depends on the local electric field [13]. The biexponential nature of the fluorescence decays observed in this study required a two-component phasor analysis involving:

- loading the dataset
- defining ROIs corresponding to individual bacteria
- defining the calibration phasor
- computing the calibrated phasor of each ROI
- determining the phasor references to be used for mixture decomposition
- computing the average lifetime of individual ROIs based on these references
- creating a color-coded map of average lifetime

The first step of loading the dataset (the example from with Fig. 3 of ref. [13] used herein can be found in the provided FigShare data repository [18]) involves a mere drag and drop of the corresponding file (FOV1.hdf5) into the main AlliGator window, resulting in the display of the intensity image (sum of all photons detected at each pixel, irrespective of their arrival time, see Fig. 4a).

**Figure 4.**
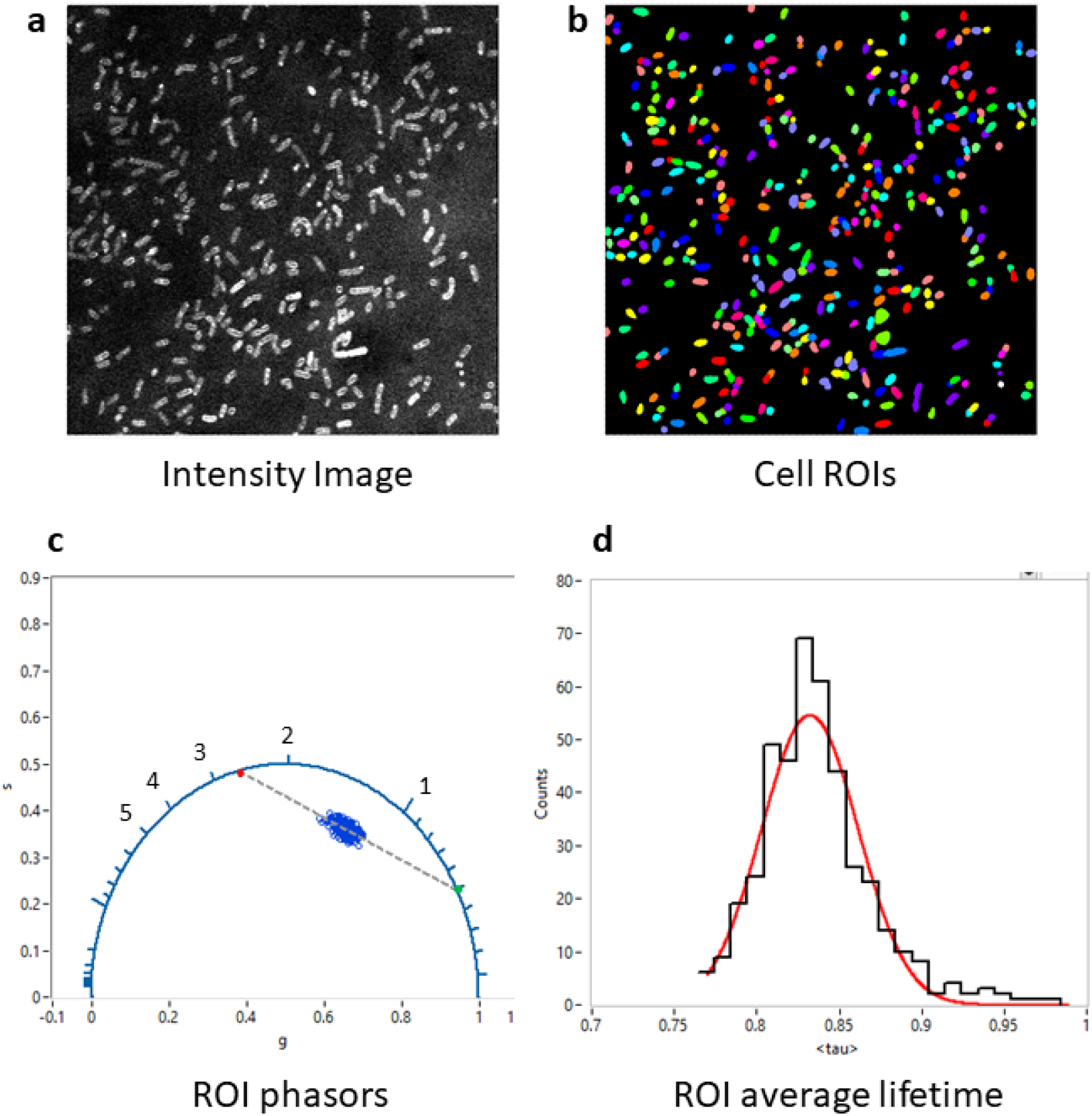
Example of phasor analysis using AlliGator. After loading the dataset (a) and defining ROIs (b), the phasor of each ROI is calculated after calibration with the appropriate instrument response function (c). A function of AlliGator’s `Phasor Graph` allows computing single-exponential *references* as the intersection of the universal circle (blue circular arc) and the major axis of the phasor scatter plot (green and red dots connected by a dashed line. These references can then be used to decompose each phasor into a linear combination of both of them, from which the amplitude-averaged lifetime *⟨τ⟩* can be computed for each phasor. The histogram in (d) is the result of a two-step analysis: first, the creation of a *⟨τ⟩* (*I*) plot and next, the calculation of the histogram of *⟨τ⟩* ‘s and its fitting with a Gaussian distribution. All images and graphs are direct export from AlliGator to the clipboard. The raw data for each plot can also be exported as an ASCII file for further analysis by third party software.

The second step consists in defining ROIs identifying each bacterial cell. In this work, we used the intensity image shown in Fig. 4a as the input to the StarDist plugin [19] of Fiji [20] to automatically identify the bacteria, and exported the corresponding label image as a TIFF file (mask FOV1.tif in [18]). This TIFF file is then loaded into AlliGator as a *mask image*, with the different ROIs automatically highlighted by different colors in Fig. 4b.

After specifying the correct phasor frequency *f* (*f* = 1*/T* = 78.02 MHz, where *T* is the laser period), and computing the phasor of the instrument’s instrument response function (IRF decay.txt), the IRF phasor was set as the calibration phasor (phasor associated with lifetime *τ* = 0) using the AlliGator *Phasor Graph*’s right-click menu item `Use Current Phasor as Calibration.`

Once phasor calibration is defined and activated by toggling the *Calibration Type* in the Phasor Graph panel to `Single Phasor`, the `Non-Interactive (Fast)` script in the `Analysis: FLI Dataset: Multiple ROIs Analysis: All ROIs Phasor Analysis` menu computes the phasors of each ROI (whose pixel signal are summed up in each ROI) and plots them as a single scatterplot in the Phasor Graph panel (Fig. 4c).

This allows defining single-exponential components describing the set of similar cells using an AlliGator-specific tool, which determines the major axis of inertia of the phasor scatterplot (see section 2.3 and Fig. 3), and determines its intersections with the universal circle (indicated as green and red dots in Fig. 4c).

The next step consists in using these references to compute the amplitudeaveraged lifetime *⟨τ⟩* _*a*_ of each ROI [13] and histogram their values to obtain the mean and standard deviation of the observed lifetime in the bacterial population (Fig. 4d).

Finally, a color-coded map of average lifetimes computed using these same references can be created as illustrated in Fig. 1. Conversion of these lifetimes into membrane potential values can then be obtained using an experimental *V* (*τ*) calibration curve.

The complete output of this analysis can be found in the Analysis folder of the FigShare repository associated with this article [18].

### 4. Impact

AlliGator is designed to analyze data in terms of single- or double-exponential decays, either by NLSF, MLE or phasor analysis. It can be used for instance to analyze mixtures of fluorophores characterized by different lifetimes, and in particular, mixture of fluorescence resonance energy transfer (FRET) donor-acceptor pairs with donor only species, as demonstrated in a series of recent works [8, 11, 13, 14, 21, 22]. It allows performing pixel-wise, ROI-wise, multi-ROIs or full dataset analysis seamlessly, with the simple selection of a menu item.

While it is fairly complete, it is in no way exhaustive. To compensate for its current limitation in terms of publication-quality graph outputs, all results can be exported in text format for further editing or processing with 3rd party software. Likewise, users with Python programming expertise can easily add functionalities in the form of plugins, which are then automatically added to the software’s main menu or to specific objects (graph, image) in the software.

AlliGator has been developed over more than a decade to address specific research needs in topics as diverse as *in vivo* FRET studies in live mice [8, 11] or membrane potential measurements in bacteria [13], using techniques as different as raster-scanning confocal microscopy [13] or wide-field single-photon avalanche diode cameras [11, 14]. These articles have garnered more than 200 citations at the time of this writing (Google Scholar). It has been used in many other articles as the tool of reference to which to compare alternative methods (e.g. deep learning) for FLI data analysis.

The software goes beyond the usual features offered by FLI analysis software, each function being documented in an extensive online manual created using open source tools (Sphinx [23] and ReadTheDocs [24]) and under version control using Git. It is designed with reproducibility in mind, all intermediate results and analysis steps being logged in a rich text format notebook window, to which the user can add and format whatever information they need.

Its ease of use and performance has made it the tool of choice for our laboratory and our collaborators (about a dozen users), who are using it on a daily basis to analyze data [8, 9, 10, 12, 13, 14, 21, 22, 25, 26, 27, 28, 29, 30, 31] and as a benchmark against other methods such as newly developed deep learning algorithms [22].

Up until this source code release, AlliGator had been freely available as a standalone executable installer on GitHub [17]. While GitHub does not provide long term visitor counts, the repository is regularly visited as indicated by the *Traffic* tab of the repository page.

## 5. Conclusions

AlliGator offers a powerful platform for FLI data analysis by NLSF or MLE bi-exponential fits and the phasor approach. The release of its source code provides the opportunity for users to check algorithmic implementation, as well as potentially suggest performance improvements or additional functionalities, either to facilitate implementation of Python plugins, or by contributing new LabVIEW code to the software. To our knowledge, it is one of the rare large size scientific software projects using this unique programming language to be released as an open source project, and with an extensive online manual and documentation.

## Acknowledgments

This work was supported in part by Human Frontier Science Program Grant RGP0061/2015, US National Institute of Health Grants R01 GM095904 & R01 CA250636, University of California CRCC Grant CRR-18-523872, US Department of Energy Grant DE-SC0020338 & DE-SC0023184, the European Research Council under European Union’s Horizon 2020 research and innovation program grant agreement No. 669941 and by the Partner University Fund, a program of the French American Culture Exchange.

It is my pleasure to acknowledge critical feedback, bug reports and feature suggestions from Rinat Ankri, Debjit Roy, Kiran Bharadwaj (Weiss lab, UCLA), Sez-Jen Chen, Jason Smith, Vikas Pandey, Nanxue Yuan, Ismail Erbas, Saif Ragab (Intes lab, RPI) and Catherine Sherry (Barroso lab, AMC).

## Appendix A. Other Fluorescence Lifetime Imaging analysis tools

Table A.3, inspired by a similar list compiled on the PhasorPy documentation page, lists some free FLI analysis software packages or plugins. The information associated with each software was collated at the time of writing this article. Code limited to uncommon file formats (and therefore not easily testable) is not discussed. Commercial software with no downloadable trial version are not discussed either.

## Appendix B. AlliGator VI Hierarchy

Fig. B.5 shows a tree view of the relationship between virtual instruments (VI) libraries (larger rectangles with icons) comprising AlliGator’s Source Code. The figure can be zoomed in for a closer look at the respective relationship between libraries and some individual VIs. A full resolution and navigable tree view can be obtained using the built-in LabVIEW tool `View:VI Hierarchy` with the `AlliGator VI Collection.vi VI` opened.

**Table A3:** OS: operating system* (W: Windows, M: macOS, L: linux, A: all), exe: standalone executable, Open: Open Source, License: Open Source License (U: unknown, BSD-3: Berkeley Software Distribution 3-clause), Language: programming language, Phasor: phasor analysis (n: n-component phasor analysis), NLSF: NLSF analysis (n: n-component NLSF analysis), MLE: MLE analysis (n: n-component MLE analysis), Docs: documentation, Year: last release date. (*) Note that Windows-only software such as AlliGator can be run in other operating systems (e.g macOS or Linux) using a Windows virtual machine (Virtual Box is an example of free and open source virtualization software that can be used for that purpose). The Berkeley Software Distribution (BSD) 3-clause license used for AlliGator imposes minimal restrictions on the use and distribution of the licensed software, allowing reproduction and modification of the source code as well as of binaries (standalone executable) as long as the original copyright holder and disclaimer are mentioned in the release.

| Name | OS | exe | Open | License | Language | Phasor | NLSF | Docs | Year |
| --- | --- | --- | --- | --- | --- | --- | --- | --- | --- |
| PhasorPy | A | Yes | Yes | MIT | Python | 4 | No | Yes | 2025 |
| SimFCS | W | Yes | No | NA | NA | 4 | No | Yes | 2021 |
| ImageJ Time Gated Phasor | A | No | Yes | U | Java | 1 | No | Yes | 2013 |
| Napari Phasor Plotter | A | No | Yes | BSD-3 | Python | 1 | No | Yes | 2025 |
| FLUTE | A | Yes | Yes | BSD-3 | Python | 1 | No | Yes | 2023 |
| GSLab | A | No | Yes | MIT | Python | 1 | No | Yes | 2025 |
| BrightEye | A | No | Yes | CC-BY-NC 4.0 | Python | 1 | No | Yes | 2022 |
| FLIM_Tools | A | No | Yes | GPL v2 | Python | 1 | No | Yes | 2022 |
| FLIMLib/FLIMJ | A | Yes | Yes | GPL v3 | C/Java | No/Yes | 3 | Yes | 2025 |
| PAM | A | Yes | Yes | GPL v3 | MATLAB | Yes | 3 | Yes | 2024 |
| AlliGator | W | Yes | Yes | BSD-3 | LabVIEW<br>Python | Yes | 2 | Yes | 2025 |

**Figure B5:**
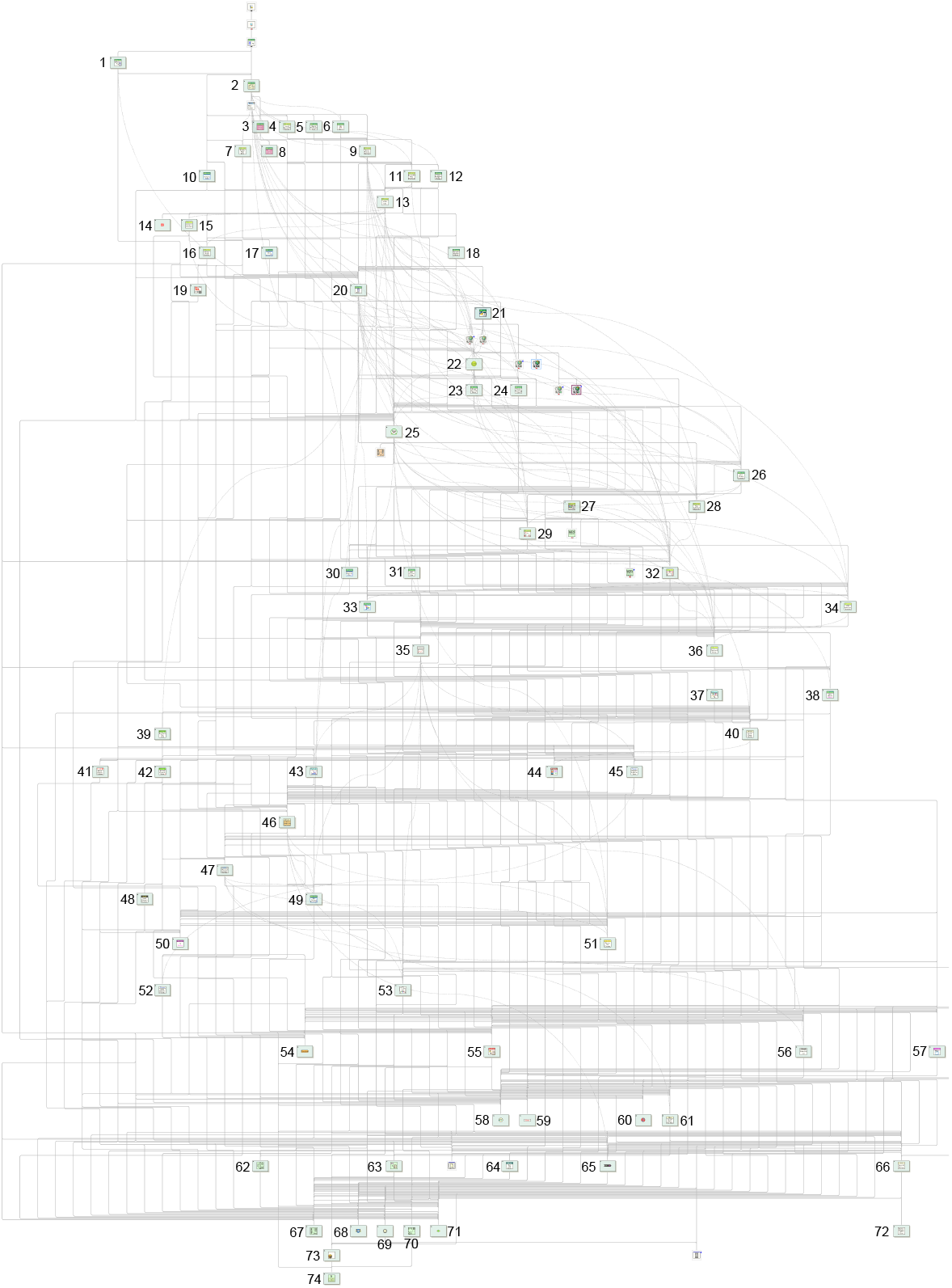
Overview of the relationship between VI libraries comprising AlliGator. The libraries are identified by numbers in the figure, as follows. *AlliGator-specific libraries:* Accumulated Dataset (12), Action Engine (2), Dataset Information Window (10), Debug (6), Decay Analysis (9), Decay Fit Parameter Map (11), Decay Statistics (18), Dual-Channel Datasets (15), Files (13), Files Tests (8), Fit Method Benchmark (49), Global Decay Fit (5), Globals, Variables & Constants (20), Graphs (23), GUI (22), HDF5 (16), Image Profile Window (7), Intensity Corrections (30), Internal Variables (27), Lifetime (31), Local Decay Window (4), Notebook (38), Phasor Harmonics (1), Phasor Calibration (28), Phasor Graph (26), Phasor Plot (32), Phasor Plot Color Map (33), Phasor Ratio (29), Python Plugins (21), ROIs (34), Scripts (24), Settings (25), Shot Noise Influence on Average Lifetime (17), Source Image (36), Test Suite (3). *Non-AlliGator-specific libraries:* Arrays (46), Becker & Hickl Files (48), Boolean (14), Buttons (72), Comparison (54), Error (71), Files (61), Fits (51), Formula (57), Graphs (56), GUI (66), Histograms (37), Histogram Window (53), Image (40), Math (47), Menu (64), Notebook (55), openg array (67), openg error (74), openg file (62), openg string (70), openg variant (73), openg variant configuration Palette (44), Phasor (45), Phasor Explorer (52), PicoQuant (39), Plot Editor (43), PTU Files (42), Rich Text Box (59), Sound (60), String (65), SwissSPAD (19), SwissSPAD Live (41), Time (68), Variant (5), Utilities (68), Variant to Data (58), XY Graph Add-Ons (35).

Libraries and VIs specific to AlliGator are described in the `Source Code Documentation` found in the `src-docs` folder of the GitHub repository [32]. Ancillary libraries’ VIs can be found in corresponding folders of the distribution. Third party libraries are listed in the Readme file accompanying the distribution.

